# Chimeric Reference Panels for Genomic Imputation

**DOI:** 10.1101/2025.04.22.648973

**Authors:** Meikun Zhou, Maddie E James, Jan Engelstädter, Daniel Ortiz-Barrientos

## Abstract

Despite transformative advances in genomic technologies, missing data remains a fundamental constraint that limits the full potential of genomic research across biological systems. Genotype imputation offers a remedy by inferring unobserved genotypes from observed data. However, conventional methods typically rely on external reference panels constructed from complete genome sequences of hundreds of individuals, a costly approach largely inaccessible for non-model organisms. Moreover, these methods generally overlook novel genomic positions not captured in existing panels. To overcome these limitations, we developed Retriever, a framework that bypasses the need for external reference panels. Retriever constructs a chimeric reference panel directly from the target samples using a sliding-window approach to identify and retrieve genomic partitions with complete data. By exploiting the complementary distribution of missing data across samples, Retriever assembles a panel that preserves local patterns of linkage disequilibrium and captures novel variants. Integrated with Beagle, Retriever achieves genotype imputation with accuracy consistently exceeding 95% across diverse datasets, including plants, animals, and fungi. By eliminating the need for costly external panels, Retriever offers an accessible, cost-effective solution that broadens the application of sophisticated genomic analyses across species.

## Introduction

Genomic research has transformed our understanding of biological systems, from elucidating phenotype-genotype associations (Visscher *et al*. 2017) to uncovering mechanisms of adaptation (Barrett and Hoekstra 2011). This transformation is fueled by technological advances: next-generation sequencing now generates over 100 gigabases per run (Goodwin *et al*. 2016), while long-read technologies produce continuous sequences exceeding 100 kilobases (Logsdon *et al*. 2020). Despite these advances, missing genomic data persists as a fundamental constraint. Datasets routinely contain substantial gaps of genotypes. This can produce inaccurate population diversity and differentiation statistics (Schmidt *et al*. 2021; Bailey *et al*. 2025), distort the construction of linkage maps (Hackett and Broadfoot 2003), and reduce power and increase false discovery rates for studies of rare variants (Auer *et al*. 2013).

Genotype imputation addresses these challenges by inferring unobserved genotypes through linkage disequilibrium patterns and haplotype structure (Li *et al*. 2009). Methods employing hidden Markov models, such as Beagle (Browning and Browning 2016), IMPUTE2 (Howie *et al*. 2009) and MaCH (Li *et al*. 2010) demonstrate exceptional effectiveness. The impact is substantial: human genomics studies have expanded from thousands of directly genotyped variants to millions of imputed variants in projects like the UK Biobank (Bycroft *et al*. 2018). Imputation has significantly enhanced genomic prediction accuracy for economically important agricultural traits (Gorjanc *et al*. 2017), enabled more cost-effective breeding strategies (Tsai *et al*. 2017), and shows growing potential to strengthen conservation genomics efforts (Theissinger *et al*. 2023) and ecological research (TaŞ *et al*. 2021).

However, these sophisticated imputation frameworks such as Beagle, IMPUTE2 and MaCH share a critical limitation: dependence on external reference panels comprising hundreds of fully genotyped individuals that represent the genetic diversity of the target population (Das *et al*. 2016; Phocas 2022a; Dekeyser *et al*. 2023a). This requirement creates a methodological divide in genomic research. While millions of human genomes have been sequenced, very few species possess reference panels suitable for conventional imputation. Panel construction demands substantial resources, specialized expertise, and infrastructure that tend to be available only for major model organisms. Additionally, conventional imputation approaches ignore and disregard any novel genomic positions in the target samples that are absent from the reference panel (Marchini and Howie 2010; Huang and Tseng 2014). This problem intensifies as sequencing technologies advance and capture novel variants with potential functional importance.

Here we introduce Retriever, a framework designed to make conventional imputation algorithms —that typically require an external reference panel— accessible to non-model organisms. Retriever creates a chimeric reference panel directly from the target dataset using a sliding-window approach that identifies genomic partitions with complete data across multiple samples. By maintaining local patterns of linkage disequilibrium within these windows, the algorithm preserves the haplotype structure critical for accurate imputation. The chimeric reference panel effectively reconfigures the patchy landscape of missing data by leveraging the observation that, while any single sample may have missing genotypes, the collective dataset often contains complete information at each position across different subsets of individuals.

We use rigorous benchmarking to demonstrate that Retriever, when used in combination with Beagle, maintains >95% imputation accuracy across allele frequency spectra in datasets without a genome reference panel. Comparative analyses against both reference-based and reference-free methods confirm that our chimeric panel strategy combines the accessibility of reference-free approaches with the precision of reference-based algorithms while minimizing disruption to underlying linkage patterns. Retriever thus makes previously inaccessible imputation algorithms available for non-model organisms, enabling researchers to extract maximum information from genomic resources and potentially reveal novel patterns otherwise obscured by missing data.

## Materials and methods

### Input files and requirements

Retriever operates on Variant Call Format (VCF) files v4.0 or higher. We use the term ‘target samples’ to refer to the collection of individuals containing missing genotypes that require imputation. The algorithm assumes that observed genotypes in the VCF file are error-free. Therefore, users must have previously implemented comprehensive quality control procedures, including filtering based on quality metrics and read depth (as outlined by (O’Leary *et al*. 2018). This preprocessing should include the removal of multiallelic sites, which can complicate haplotype inference, and the elimination of positions with greater than 20% missing data.

As Retriever uses a sliding-window approach, it should be executed on a per-chromosome or contig basis. Importantly, Retriever must be implemented on cohorts of genetically similar samples, such as individuals within a population, ecotype, or species, as substantial genetic divergence is expected to compromise the underlying assumption of shared haplotype structures. GVCF files, which contain additional reference blocks for non-variant regions, are not encouraged for use in Retriever. The implementation of Retriever uses established Python packages for processing and manipulating VCF files. A list of external dependencies is documented in the project’s Github repository (https://github.com/uqmzhou8/Retriever.git), and a YAML configuration file is available to facilitate the seamless installation of these dependencies.

### Assembly of the chimeric reference panel

The central innovation of Retriever is the construction of a chimeric reference panel from the target data itself, thereby eliminating the need for an external reference panel (Fig 1). This chimeric assembly process identifies and retrieves substantial genotype information across the genomic landscape of the population. To construct the chimeric reference panel, Retriever first parses the VCF file and implements a non-overlapping sliding-window approach with a user-defined size (default: 1000bp) that sequentially scans all genomic positions within the target samples. For each window, the algorithm identifies sets of complete genotype partitions— groups of individuals with no missing genotypes within the current window— and stores these partitions in separate data structures, or “buckets.” Prior to storage, the algorithm verifies that the number of individuals with complete genotype data in the window meets the user-defined minimum number of chimeric individuals to be generated. If this requirement is not satisfied, the window shrinks by 1bp increments until complete partitions with the minimum number of individuals are identified. Should this condition remain unsatisfied despite window contraction, potentially due to inadequate filtering of the initial dataset, the algorithm prematurely terminates and produces an error message.

**Figure 1.**
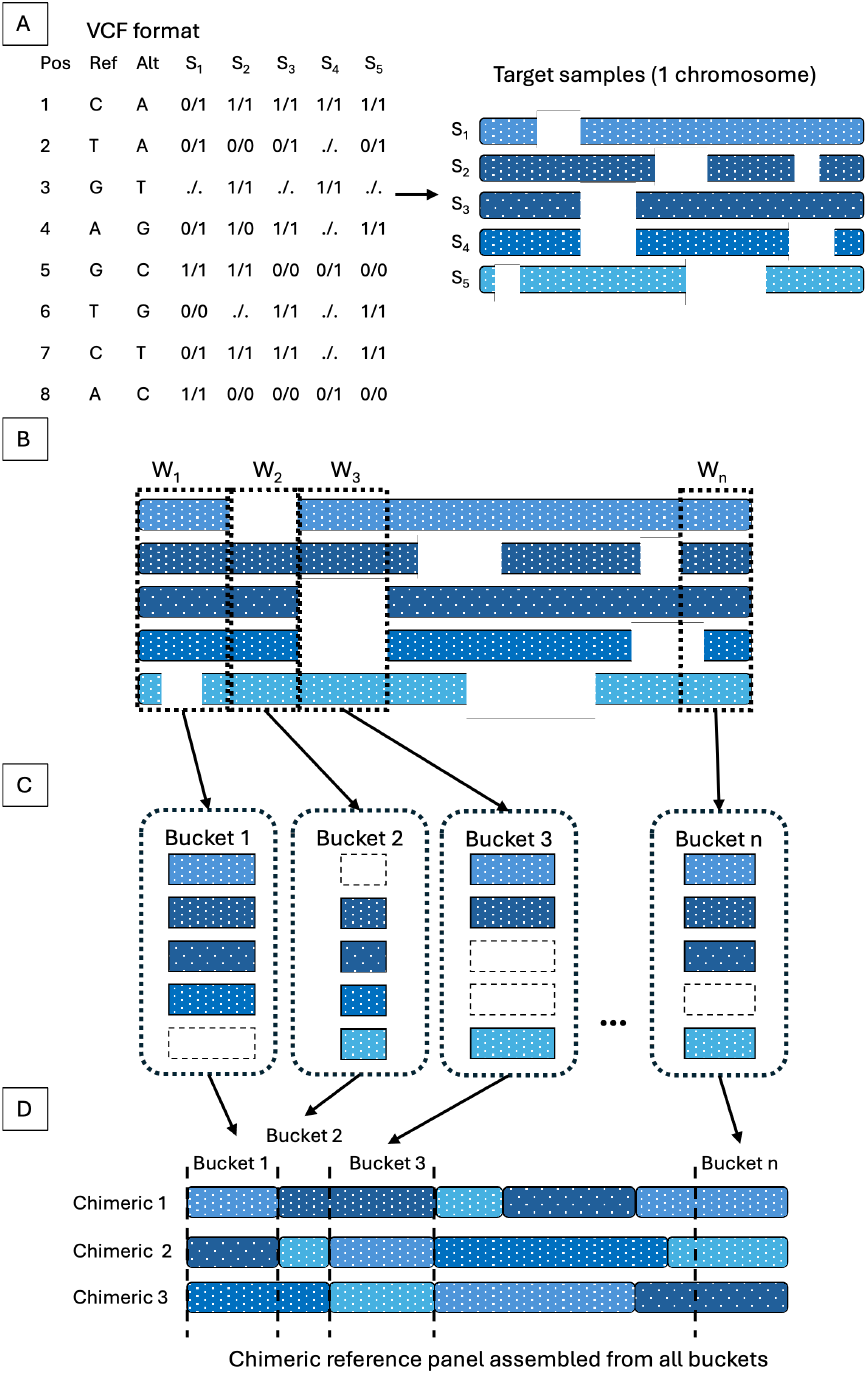
Steps for assembly of Retriever’s chimeric reference panel. (A) A VCF file containing the diploid target samples (S_1_, S_2_ … S_5_) from a single population, ecotype or species, aligned to a reference genome, is read by Retriever. Individual genomes of five target samples are represented by blue bars and gaps represent missing genotypes. (B) A non-overlapping sliding-window (W) moves across the genome, selecting complete partitions -genotypes without missing data- to be stored into buckets. If the partition of complete genotypes is less than the user-defined chimeric reference panel size, the window shrinks until the required number of partitions is reached. (C) Partitions of complete genotypes are stored in individual buckets with non-overlapping genomic positions. (D) Within each bucket, individuals are randomly selected to assemble the chimeric reference panel in contiguous genomic order. In this example, the user-defined size for the chimeric reference panel is three, so three individuals are randomly selected per bucket.

As the sliding-window progresses through the genome, the window size is reset at each new position. Upon completing the genome scan, the algorithm assembles the chimeric reference panel by chronologically concatenating the genotype partitions stored in the respective buckets. In cases where a bucket contains more individuals than the requested chimeric reference size, individuals are randomly selected to use in the final chimeric reference panel. This assembly procedure yields a chimeric reference panel comprising individuals with contiguous, non-missing genotype data, compatible with most established imputation algorithms that require external reference panels.

### Comparison of chimeric versus external reference panels

We validated the imputation accuracy using Retriever’s chimeric reference panel against that using an external reference panel of the same size, using a well-established imputation software as a benchmark for the highest achievable imputation accuracy. We selected Beagle4.1 (beagle.27Jan18.7e1.jar) as the imputation engine due to its widespread adoption and consistently high performance reported in literature (Das *et al*. 2016). In addition, Beagle4 only requires an external reference panel to initiate the imputation process. In contrast, other imputation software, such as EagleImp (Wienbrandt and Ellinghaus 2022), require more resources, such as a genetic map. These requirements can be difficult to obtain, particularly for laboratories working on non-model organisms. For instance, a forced initiation of EagleImp without a genetic map can result in an early termination or simply an error prompt upon running the program, making it unsuitable for use with Retriever’s chimeric reference panel.

Each analysis was replicated three times by randomly masking different locations of the target samples to assess consistency and derive statistical estimates. All analyses presented in this study were executed on the National Computational Infrastructure in Australia. The assembly of the chimeric reference panel was performed using a 24-core Intel Xeon Platinum 8274 (Cascade Lake), while imputation with Beagle4 used a 24-core Intel Xeon Platinum 8268 (Cascade Lake) processor. The imputation process used 96 threads with default parameters of Beagle4 (Browning and Browning 2016) consistently across all analyses to ensure uniformity in our methods.

#### Sample selection

To validate imputation accuracy and demonstrate Retriever’s broad applicability, we used data from the hu man (*Homo sapiens*) 1K Genome Project (1000 Genomes Project Consortium *et al*. 2015) alongside genomic datasets from diverse organisms, including the plant *Arabidopsis thaliana*, the animal *Gallus gallus*, and the fungus *Saccharomyces cerevisiae*. The 1K Genomes Project was chosen because of its high quality and availability of a large number of samples suitable for constructing an external reference panel for Beagle4. It is also a widely used benchmark in the field and has been employed in the validation of various imputation tools (Delaneau *et al*. 2008; Browning *et al*. 2018; Wienbrandt and Ellinghaus 2022).

The individuals in the current study were chosen to be of African origin as they have the highest nucleotide diversity among all the human populations (Yasukochi *et al*. 2019). This choice allowed us to test Retriever’s performance under a conservative scenario that represents a challenging condition for imputation, thereby providing a robust assessment of Retriever’s capabilities. From the 661 individuals of African ancestry, we randomly selected 200 individuals as target samples for imputation. From the remaining pool, 50 individuals were randomly chosen to construct the external reference panel. This external reference panel serves as a baseline for Beagle4, representing the highest achievable imputation accuracy for a panel of similar size. In contrast, the chimeric reference panel of Retriever is assembled from the target samples. We used chromosome 1 for analysis as it is the largest chromosome and reduces computational complexity compared to analyzing the entire genome.

#### Data masking

We simulated missing genomic data by randomly masking genotypes across all individuals and genomic positions to create completely missing at random data. Both alleles at each diploid site were masked to ensure there was no data leakage for downstream analysis as the presence of one allele that remains unmasked in a heterozygous site may potentially bias the probability of the identity of the other allele. Additionally, to examine how well the chimeric reference panel captures novel variants in the target samples, we removed random genomic positions from the external reference panel while keeping them in the target samples. This simulates real-world scenarios where target samples contain novel variants absent from pre-existing reference panels —a common occurrence when newer sequencing technologies or higher coverage reveal variants missed during external panel assembly.

#### Parameter optimization

We performed several tests to understand the influence of chimeric panel size on imputation accuracies. Theoretically, larger reference panels should yield better accuracy. However, the maximum size of the panel is constrained by the dataset’s filtering thresholds, as the chimeric assembly selects the most complete haplotypes within the sliding-window. Therefore, the maximum achievable size of the chimeric reference panel will depend on the number of individuals with complete data at the genomic position with the highest proportion of missing genotypes. Using the human dataset, we tested a range of panel sizes (from 10 to 100 individuals) across various masking proportions (1% to 30%) to identify the optimal balance between these competing factors. The maximum achievable size of the chimeric reference panel is inherently constrained by the dataset’s missing data patterns, as the algorithm requires complete genotypes within each sliding-window. Therefore, we evaluated how different panel size requirements affected both window size dynamics and downstream imputation accuracy.

#### Statistical framework for accuracy assessment

The imputed results obtained from the chimeric and external panels were compared to the original unmasked samples to determine the average imputation accuracy of all individuals at each position. To evaluate the imputation accuracy of each panel in relation to the minor allele count, an average accuracy from all three replicates was binned by the minor allele count associated with each position. The minor allele count was derived from the original dataset using BCFtools v1.9 (Danecek *et al*. 2021). For each genomic position, we calculated accuracy as the proportion of correctly imputed genotypes among all masked genotypes. Our method employs a stringent matching criterion: a heterozygous genotype (0/1) must be imputed exactly as 0/1 to be considered correct; an imputation of 1/0, though functionally equivalent, is counted as incorrect. This approach differs from conventional methods that often encode genotypes numerically (e.g., homozygous as 1 or 3, heterozygous as 2), which would treat 0/1 and 1/0 as identical. As mentioned, we accounted for the relationship between imputation accuracy and allele count by stratifying positions by their minor allele count (MAC).

#### Scaling of Retriever

We evaluated the computational resources (wall time) required for constructing the chimeric reference panel with varying numbers of individuals from the human dataset outlined above and different proportions of masked (missing) data. Retriever was designed to process the genome using a single continuous sliding-window from start to end across all genomic positions within a contig. While this approach increases computational time compared to a partitioned approach, it preserves the integrity of linkage disequilibrium patterns in the resulting chimeric reference panel. Partitioning each chromosome into separate segments for parallel processing would create artificial boundaries between adjacent regions, potentially fragmenting haploblocks that span these boundaries. This fragmentation would increase the likelihood of combining genotypes from different individuals at these boundaries, thereby disrupting important linkage disequilibrium patterns. The single-window approach, therefore, maintains biological accuracy while requiring modest computational resources (just 1 CPU).

### Evaluation of non-human organisms

We further expanded our validation to additional species that lack the resources to assemble an external reference panel and are, therefore, unable to use conventional genomic imputation software. To achieve this, we sourced publicly available datasets and selected organisms from various taxonomic kingdoms, including plants, animals, and fungi. For plants, we used *Arabidopsis thaliana* sequence data from the Weigel laboratory at the Max Planck Institute for Developmental Biology (Consortium 2016). From this genomic dataset consisting of 1135 individuals, 200 individuals were randomly selected for the target samples. The animal kingdom was represented by 188 individuals of chicken *Gallus gallus* data (Cho *et al*. 2022). Fungi were represented by a yeast *Saccharomyces cerevisiae* dataset of 165 individuals (Sardi *et al*. 2018). As with the human dataset, masking was applied to each set of target samples, and only chromosome 1 was used for analysis. Using these methodological approaches, we evaluated the performance of Retriever across diverse datasets and compared it with conventional imputation strategies to assess accuracy, computational efficiency, and applicability across taxa.

## Results

### Influence of chimeric reference panel size on imputation accuracy

We first examined how the size of the chimeric reference panel affects imputation accuracy across various proportions of data masking using the human dataset (Fig. 2). As expected, higher masking proportions resulted in modest reductions in imputation accuracy. However, the relationship between panel size and accuracy did not follow a simple monotonic pattern. Instead, we observed an inflection point in performance, with optimal panel sizes ranging from 25 to 50 individuals from a target sample of 200 individuals (12.5-25% of the total sample size). This pattern remained consistent across all masking levels (1-30%), suggesting that the observed optimum represents an intrinsic feature of the chimeric reference construction process rather than an artifact of data sparsity. This inflection point likely represents a fundamental trade-off in the chimeric panel construction process. As the required panel size increases, the algorithm must identify more individuals with complete genotypes within each window, often necessitating window size reduction. Initially, increasing panel size improves performance by incorporating greater haplotype diversity. However, beyond the optimal range, the benefits of additional diversity are offset by increasing fragmentation of linkage disequilibrium patterns as windows become smaller and more numerous. The consistency of this pattern across all masking levels (1-30%) suggests that this represents an intrinsic property of the methodology rather than an artifact of data sparsity.

**Figure 2.**
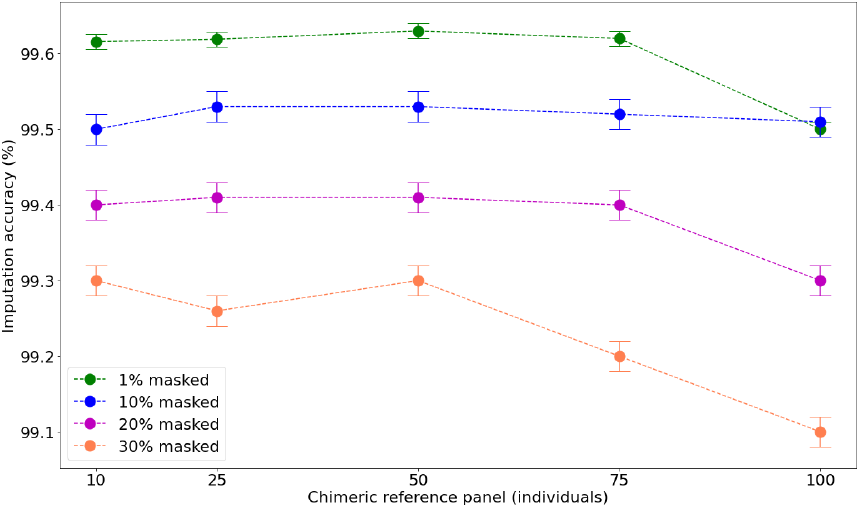
Impact of Retriever’s chimeric reference panel size on imputation accuracy. Imputation accuracy for *Homo sapiens* data obtained from the 1K Genomes Project, imputed in Beagle4. The chimeric reference panel size ranges from 10 to 100 individuals, selected from 200 individuals in the target sample. Each line represents different missing data proportions that vary from 1% to 30% masked. Points represent the mean *±* standard error from three independent replicates.

### Imputation accuracy comparison with an external reference panel

We compared the imputation performance of Retriever’s chimeric reference panel composed of 50 individuals against an external reference panel of equivalent size using the human dataset (Fig. 3). The comparison was stratified by minor allele count to evaluate performance across the allele frequency spectrum. Both panels exhibited comparable accuracy profiles, with median accuracies exceeding 0.96 at low masking levels (1%, Fig. 3A) and 0.93 even at high masking levels (30%, Fig. 3B).

**Figure 3.**
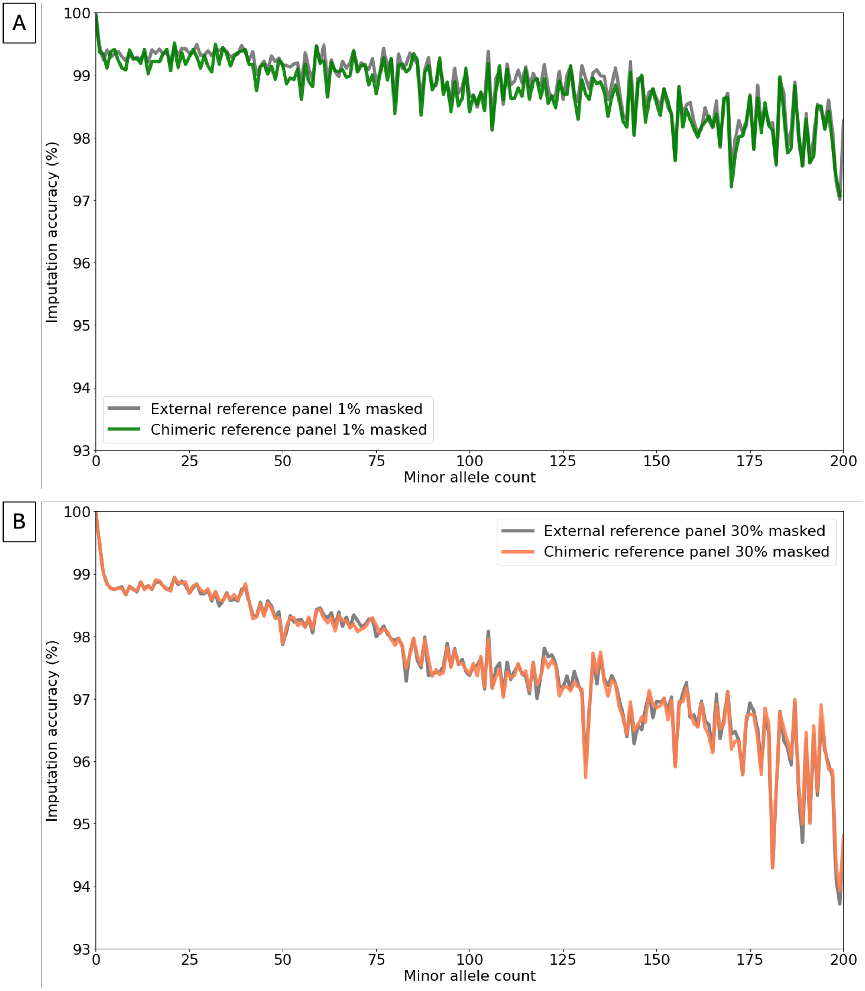
Effect of minor allele count on imputation accuracy. *Homo sapiens* data obtained from the 1K Genomes Project (2*N* = 400, *π* = 0.027-0.034 (Yasukochi *et al*. 2019)) were masked at 1% (A) and 30% (B) and imputed with Beagle4 using either an external reference panel, or Retriever’s chimeric reference panel, both with a panel size of 50 individuals. Imputed accuracies are binned according to the minor allele count from the original unmasked master file.

### Imputation of novel variants absent from external reference panels

A key limitation of conventional imputation approaches is their inability to impute genomic positions absent from the reference panel. We quantified this effect by systematically removing positions from the external reference panel while retaining them in the target samples (Fig. 4). As expected, imputation accuracy using the external reference panel dramatically declined as the proportion of removed positions increased, with accuracy falling to below 80% when 20% of positions were absent from the reference panel. In contrast, Retriever’s chimeric reference panel maintained close to perfect imputation accuracy. This fundamental advantage stems directly from Retriever’s methodology. By constructing the reference panel from the target samples themselves, all variants present in the dataset—including those that would be novel relative to an external panel—are represented in the chimeric reference panel. The algorithm’s sliding-window approach ensures that each genomic position is captured within at least one window, allowing the imputation engine to leverage local haplotype patterns even for variants that conventional approaches would traditionally discard. This capability becomes increasingly important as sequencing technologies continue to advance, revealing previously undetected genetic variation that might hold biological significance.

**Figure 4.**
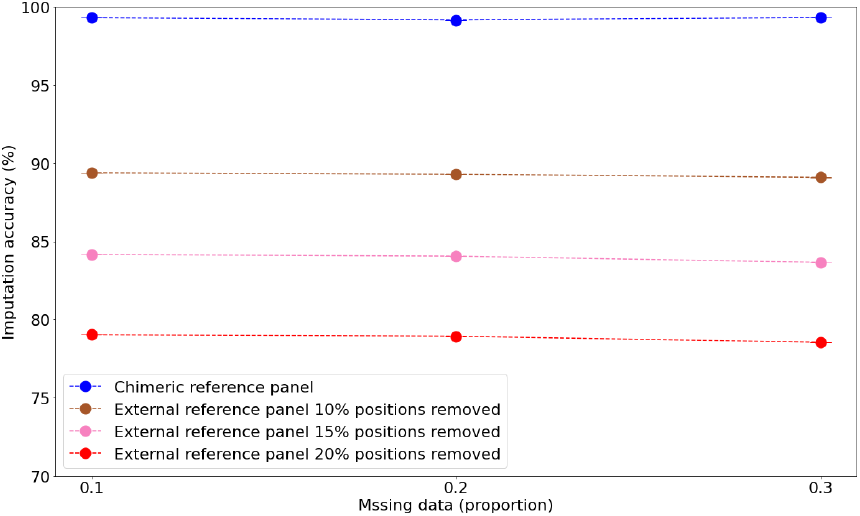
Beagle4’s performance on partially observed data. Target samples are 200 *Homo sapiens* individuals with 10% genotypes masked. Imputation of the target samples were compared using either a chimeric reference panel assembled with Retriever or an external reference panel that had random positions removed across all samples (10%, 15% and 20% positions removed). Points represent the mean*±* standard error from three independent replicates.

### Computational scaling and resource utilization

We evaluated the computational resources required for chimeric reference panel assembly under varying conditions of sample size and data masking (Fig. 5). Computational time (wall time) remained relatively stable when missing data (masked positions) was low (1-10%) but increased substantially when missing data exceeded 20%. Importantly, for the optimal panel size range identified earlier (25-50 individuals from 200 samples), chimeric assembly required approximately 28 hours even with 30% missing data. These computation times represent a modest investment for most research groups. Running on a standard high-performance computing environment with a single CPU core, the entire process can typically be completed overnight for most chromosomes. The relationship between missing data proportion and computational time suggests that preprocessing steps to remove positions with excessive missingness (>20%) can significantly improve efficiency. For context, these computational requirements are negligible compared to the resources needed to generate an external reference panel, which typically involves years of sequencing efforts, substantial financial investment, and collaborative networks—resources largely unavailable for non-model organism research. In general, Retriever should be accessible to research groups with limited resources, widening access to high-quality imputation capabilities.

**Figure 5.**
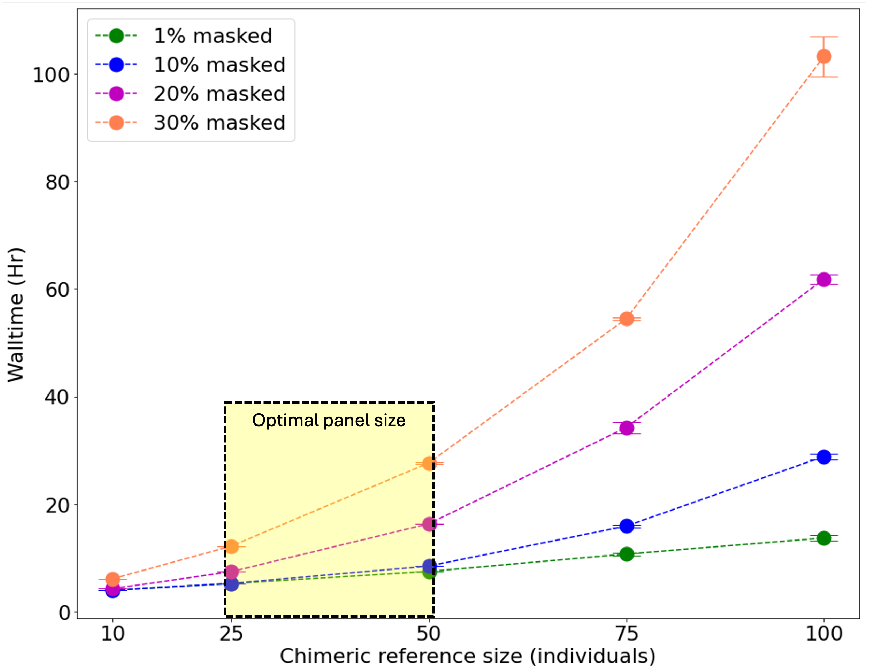
Wall time plot showing the impact of varying proportions of masked data on chimeric reference assembly. Longer wall times indicate increased computational demand. Yellow box represents the optimal chimeric reference panel size, obtained from Figure 2. Points represent the mean*±* standard error from three independent replicates.

### Cross-species validation of imputation accuracy

To demonstrate Retriever’s broad applicability, we extended our validation to diverse model organisms spanning plants (*Arabidopsis thaliana*), animals (*Gallus gallus*), and fungi (*Saccharomyces cerevisiae*) (Fig. 6). Imputation accuracy consistently exceeded 95% across all species and levels of missing data, with minor variations likely corresponding to the underlying genetic diversity of each dataset. The *A. thaliana* dataset, derived from inbred lines with relatively homogeneous genetic structure, achieved the highest accuracy, whereas the *S. cerevisiae* dataset, representing 8 distinct populations, exhibited slightly lower but still excellent accuracy. The more genetically diverse *G. gallus* dataset, comprising 13 populations, showed the lowest accuracy, consistent with the expected negative relationship between population diversity and imputation performance.

**Figure 6.**
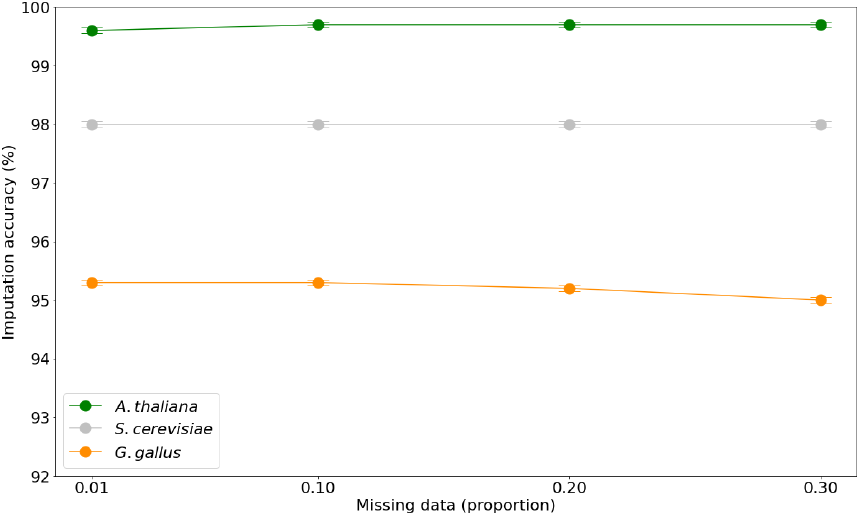
Imputation accuracy of non-human model organisms. Chimeric reference panels, of 50 individuals, were constructed with genomic data from *Arabidopsis thaliana, Gallus gallus* and *Saccharomyces cerevisiae*, and imputed in Beagle4 across different proportions of masked (missing) data. Points represent the mean *±* standard error from three independent replicates.

## Discussion

In this study, we developed Retriever, a novel framework that enables high-accuracy imputation for species lacking external reference panels. Through rigorous benchmarking, we demonstrated that Retriever’s chimeric reference panel approach maintains imputation accuracy above 95% across diverse taxonomic groups while eliminating the need for resource-intensive external reference panel construction. Here, we examine the implications of our results, explore the theoretical principles underlying Retriever’s effectiveness, and outline future directions for improvement.

### Methodological Foundations and Statistical Principles of Retriever

The fundamental innovation of Retriever lies in constructing a reference panel directly from the target data to be imputed. Our comparative analysis revealed that this chimeric reference panel performs comparably to external reference panels of equivalent size. This equivalence is remarkable, considering that external reference panels typically require extensive construction resources (Browning and Browning 2016; Sengupta *et al*. 2023), often involving dedicated sequencing efforts to ensure complete genotype information across hundreds of individuals. Both panel types showed similar performance patterns: imputation accuracy decreased as the minor allele count increased (reflecting greater genetic diversity) and as the proportion of missing genotypes increased (reducing available information about population structure).

When applying Retriever to diverse non-human organisms, we observed consistent imputation accuracy above 95% across species, with imputation accuracy correlating with population structure complexity. The *G. gallus* dataset, comprising samples from 13 distinct populations (Cho *et al*. 2022), showed the lowest accuracy, reflecting challenges in imputing across high genetic diversity. Conversely, *S. cerevisiae* (8 populations) (Sardi *et al*. 2018) and *A. thaliana* (inbred lines) (Consortium 2016) demonstrated higher accuracy (approximately 98%), benefiting from their more homogeneous genetic backgrounds. These results indicate that Retriever performs best when applied to samples with relatively low intra-sample genetic diversity, such as within a single population or closely related individuals. We, therefore, recommend using Retriever primarily on samples composed of closely related individuals or within a single population. This guideline aligns with common research practices of standard imputation software (Huang and Tseng 2014; Marino *et al*. 2022).

#### Theoretical foundations of chimeric reference panel construction

The chimeric reference panel construction takes advantage of specific properties of genomic organization and the statistical distribution of missing data. Retriever exploits the non-random patterns of linkage disequilibrium, haplotype block structure (Mochurad and Horun 2023; Dekeyser *et al*. 2023b), and recombination rate variation (Smukowski and Noor 2011) across the genome, alongside the probabilistic complementarity of sequencing coverage across multiple samples (Nunney 2001). By “probabilistic complementarity of sequencing coverage,” we refer to the statistical phenomenon whereby the patterns of missing data tend to vary stochastically across samples in next-generation sequencing datasets.

As sequencing depth varies across genomic regions due to factors such as GC content (Benjamini and Speed 2012; Aird *et al*. 2011; Huang and Knowles 2016), sequence complexity (Sims *et al*. 2014), structural features (Muyas *et al*. 2019), and stochastic sampling, missing genomic data rarely follows a completely random pattern (MCAR) (Wong *et al*. 2019). This non-random distribution creates a favorable scenario: the probability that any specific position will be missing in all samples decreases exponentially with increasing sample size. Consequently, despite substantial missing data in individual samples, the collective dataset retains information at nearly all genomic positions across different subsets of individuals. Retriever exploits this fundamental statistical property to construct complete chimeric reference panels. Our recommended preprocessing workflow removes genomic positions with more than 20% missing data prior to chimeric panel construction, effectively eliminating sites most vulnerable to comprehensive information loss across samples. Nonetheless, researchers should remain cognizant of potential biases when interpreting results from regions known to be problematic for sequencing technologies.

The chimeric reference panel construction leverages how genetic variants are naturally organized in genomes. Genetic variants are not randomly distributed (Cheung *et al*. 2011; Dong *et al*. 2021), they show patterns of association with nearby variants that reflect the population’s evolutionary history, including past recombination events and selection pressures (Edwards and Beerli 2000; Hellenthal and Stephens 2006). Retriever’s sliding-window approach seeks to preserve these important genetic patterns by identifying genomic partitions with complete genotype information. However, this preservation is contingent upon the distribution of missing data, which could break down these associations; if missing genotypes occur non-randomly and cluster in regions of strong linkage disequilibrium (LD) or at functional elements, the effectiveness of the sliding-window approach may be compromised. Our empirical validation suggests that under typical sequencing scenarios, sufficient complementarity exists to maintain the integrity of the local LD structure.

Our optimization testing revealed that chimeric panels with 25-50 individuals (about 12.5-25% of the total samples) achieved the highest accuracy. This finding makes sense when considering the tradeoffs involved. Larger panels can capture more genetic diversity but force the algorithm to use smaller window sizes to find enough complete data segments. When windows become too small, they break up the natural patterns of genetic association that help predict missing genotypes. Based on these findings, we recommend setting the chimeric reference panel size to about 25% of the total samples or 50 individuals, whichever is larger.

Retriever also takes advantage of the natural constraints on genetic variation. The combinations of genetic variants we observe are not random—they are shaped by evolution, natural selection, and population history (Akey *et al*. 2004; Zhang *et al*. 2013; Smukowski and Noor 2011). These constraints limit the possible genotypes when predicting missing data, making imputation more tractable. While rare genetic variants are generally harder to impute correctly (Lau *et al*. 2024) (they provide less information for the algorithm to work with), our chimeric reference panel method performs similarly to traditional reference panels across all variant frequencies, suggesting our approach successfully captures the underlying genetic patterns that determine which variant combinations are biologically likely.

#### Implications of stochastic sampling in chimeric panel construction

The assembly of the chimeric reference panel involves randomly selecting individuals from each “bucket” of complete genotypes within defined genomic windows. This stochastic component introduces both advantages and potential limitations. On the one hand, the randomization process prevents ascertainment biases while maintaining the fundamental property of complete genotype information within each window. However, random selection could remove rare variants, and potentially disrupt trans-window haplotype continuity when transitions occur between adjacent genomic regions. Our empirical analyses suggest that this concern has minimal practical impact on imputation accuracy. This resilience likely reflects the primacy of local LD patterns in determining imputation accuracy; that is, preserving complete haplotype information within windows appears more critical than maintaining long-range haplotype continuity across window boundaries. Nevertheless, future iterations of Retriever could implement more sophisticated sampling strategies that consider haplotype similarity at window boundaries. For instance, we could prioritize matching individuals across adjacent windows when possible. If we have sampled individuals (3,7,1,2) in one window and individuals (8,11,5,7) in the next, we could create concatenated chimeric reference sequences by connecting individuals with shared identities first (like individual 7), then completing the remaining connections (e.g., 3-8, 7-7, 1-11, and 2-5). This approach would potentially enhance trans-window haplotype continuity while maintaining the core advantages of the chimeric assembly process.

#### Advantage of capturing novel variants

A significant advantage of Retriever over conventional methods is its ability to impute novel variants absent from external reference panels. As sequencing technologies advance, newer studies often identify variants or indels not present in existing reference resources (Satam *et al*. 2023). Conventional imputation approaches that rely upon external reference panels discard these novel positions, potentially losing valuable information (Lau *et al*. 2024). By contrast, Retriever retains and processes all variants detected in the target dataset during imputation. This means population-specific variants and newly discovered polymorphisms are incorporated into the imputation framework rather than being systematically excluded. This inclusion is particularly valuable for capturing functionally relevant variation that might be population-specific or absent from established reference resources. Our analysis demonstrates that Retriever maintains significantly higher imputation accuracy compared to external reference panels when target samples contain novel genomic positions, ensuring that valuable information is preserved rather than discarded.

### Resource utilization and efficiency

The computational demands of assembling a chimeric reference panel are modest compared to the resources required for constructing an external reference panel (1000 Genomes Project Consortium *et al*. 2015; Tadaka *et al*. 2021). Even with 30% masked data, assembling an optimal chimeric panel typically takes less than 30 hours of wall time. Although this represents a significant computational period, it is a negligible cost compared to the years of sequencing efforts, substantial financial investment, and collaborative networks required to build an external reference panel—resources typically unavailable for non-model organism research.

By shifting the resource burden from wet-lab sequencing to computational analysis, Retriever provides an economical solution for genomic imputation while maintaining high accuracy. This accessibility has particular significance for conservation genomics (Fuentes-Pardo and Ruzzante 2017), biodiversity research (Escalante *et al*. 2014), and studies of neglected crop species (Liu 2011; Esposito *et al*. 2016; Ashraf *et al*. 2022), where resource limitations have historically constrained methodological options.

### Comparative advantages of Retriever over other methodologies

Contemporary genomic imputation methodologies can be categorized along a methodological spectrum, ranging from reference-based approaches to reference-free strategies, each with distinct conceptual foundations and operational constraints. Retriever occupies a strategic intermediate position, synthesizing advantageous elements from both paradigms while addressing their respective limitations.

#### Reference-based methodologies: Statistical power with resource limitations

Reference-based approaches, exemplified by established frameworks such as Beagle (Browning and Browning 2016), IMPUTE2 (Scheet and Stephens 2006), IMPUTE5 (Rubinacci *et al*. 2020), Minimac (Howie *et al*. 2012), and more recently EagleImp (Wienbrandt and Ellinghaus 2022) and Minimac4 (Das *et al*. 2016), rely on comprehensive external reference panels and sophisticated statistical models. These methods predominantly implement hidden Markov models (HMMs) or positional

Burrows-Wheeler transforms to infer unobserved genotypes through probabilistic modeling of haplotype structure. Their statistical power derives from capturing population-level patterns of linkage disequilibrium encoded within reference panels (Phocas 2022b) comprising hundreds to thousands of fully genotyped individuals.

However, reference-based approaches face substantial constraints. EagleImp and IMPUTE5, despite offering state-of-the-art accuracy for human genomics, require not only reference panels but also genetic maps and phased reference panels, creating resource requirements that restrict their applicability beyond model organisms. Even Beagle4, which shows greater flexibility, exhibits diminishing returns when reference panel size decreases below certain thresholds—a particular challenge for non-model organisms where comprehensive genomic resources remain unavailable. Moreover, these methods discard genomic positions absent from reference panels, excluding potentially significant novel variation. This limitation creates a methodological blind spot for population-specific variants, which often hold particular functional or evolutionary significance (Huang and Tseng 2014). This constraint becomes increasingly problematic as sequencing technologies advance, progressively revealing previously undetected genetic diversity.

#### Reference-free approaches: Accessibility with black-box complexity

At the opposite end of the methodological spectrum, reference-free approaches have emerged to address these accessibility limitations. Recent implementations include machine learning frameworks such as GRUD (Chi Duong *et al*. 2023), which employs gated recurrent units with adversarial training, Two-Stage DAE+RNN (Kojima *et al*. 2024) that combines denoising autoencoders with recurrent neural networks, and SOM-based imputation (Mora-Márquez *et al*. 2025) that implements self-organizing maps for pattern recognition. Other methods include GenoPop-Impute’s random forest implementation (Gurke and Mayer 2024) and FastImpute’s lightweight regression-based approach (Ge *et al*. 2024).

These methods circumvent reference panel requirements by learning genotype patterns directly from incomplete data. Their key advantage lies in accessibility, eliminating the infrastructure demands of reference panel construction. However, they introduce distinct limitations: complex hyperparameter optimization requirements, substantial computational resources for model training (often exceeding 50-100 hours for genome-wide applications), and limited biological interpretability of their underlying statistical representations (Naito and Okada 2024). Furthermore, these methods typically demonstrate 2-5% lower accuracy compared to reference-based approaches when evaluated on datasets with available reference panels (Kojima *et al*. 2024). This performance gap widens particularly for rare variants and regions with complex haplotype structures, where deep learning models struggle to capture subtle biological patterns from limited examples (Jiang *et al*. 2021).

#### Retriever’s chimeric framework: Combinining methodological strengths

Retriever strategically bridges reference- and -free and imputation paradigms through its chimeric reference panel construction. Unlike both methodological categories, Retriever uniquely maintains the established statistical frameworks of reference-based methods while eliminating external reference requirements. Further, unlike the abstract learned representations of machine learning approaches, Retriever preserves an explicit representation of haplotype structures. Retriever also imputes all genomic positions observed in the target dataset—including novel variants systematically excluded by reference-based methods, and requires minimal configuration compared to the extensive hyperparameter tuning demanded by machine learning approaches. Finally, it shows consistently high accuracy (>95%) across diverse taxa, approaching the performance of optimal reference-based methods while exceeding typical reference-free implementations by 1-3% (Chi Duong *et al*. 2023; Mora-Márquez *et al*. 2025)

Retriever’s chimeric methodology presents distinctive advantages in different contexts. For integrative studies involving both model and non-model organisms, Retriever enables methodological consistency across datasets with varying resource availability. In conservation genomics, where sample availability necessarily limits reference panel construction, Retriever can extract information from sparse data without sacrificing the needed rigor in downstream statistical analysis. For novel variant analysis in clinical genomics, Retriever’s preservation of observed genomic positions potentially captures functionally relevant variation that can be excluded by conventional approaches. The comparable performance between chimeric and external reference panels of equivalent size (Table 1) demonstrates that our approach successfully preserves essential population genetic information required for accurate imputation while substantially reducing barriers to implementation. This combination of accessibility, interpretability, and performance positions Retriever as a valuable bridge, extending genomic analysis capabilities beyond traditional model systems.

**Table 1.** Comparison of Retriever with reference-based and reference-free imputation approaches.

|  | Retriever (Chimeric Panel) | Reference-Based | Reference-Free ML |
| --- | --- | --- | --- |
| Core approach | Assembles a chimeric reference panel from target samples | Uses external reference panels comprising fully genotyped individuals | Learns patterns directly from incomplete data without reference panels |
| Accuracy | >95% across diverse taxa and masking levels | >98% for model organisms; performance degrades for non-model organisms | 93–97% depending on implementation and data characteristics |
| Novel variant handling | Preserves and imputes all genomic positions in target samples | Cannot impute positions absent from reference panel; novel variants discarded | Variable capability; some may impute genotypes in novel regions |
| Resource requirements | Moderate computational resources; no external panel needed | External panel requiring extensive sequencing efforts across hundreds of individuals | Substantial computational resources for model training; no external panel needed |
| Biological interpretability | Preserves explicit haplotype structures and linkage disequilibrium patterns | Direct modeling of population genetic parameters | Abstract learned representations without direct population genetic interpretation |
| Primary application | Non-model organisms lacking reference resources | Model organisms with established external reference panels | Non-model organisms lacking reference resources |
| Implementation complexity | Moderate; requires basic configuration of window parameters | High; dependent on reference panel availability and quality | High; requires expertise in machine learning methodologies and hyperparameter optimization |

### Future directions

While Retriever currently demonstrates strong performance across diverse taxa, several promising directions can enhance its methodology. Currently, Retriever assembles the chimeric reference panel using random selection of complete genotype partitions, but future versions could implement a more sophisticated partition ranking system. By prioritizing genomic partitions based on their likelihood of containing rare alleles or evolutionarily significant variants, Retriever could enhance imputation of low-frequency genetic variation while increasing overall panel information content. Integration with machine learning approaches represents another promising direction. Incorporating neural network architectures such as attention-based models or graph neural networks to learn complex patterns from the chimeric reference panel could potentially improve imputation in regions with limited linkage disequilibrium or complex structural variation. This hybrid approach would combine the biological interpretability of reference-based methods with the pattern recognition capabilities of machine learning.

These methodological enhancements would expand Retriever’s applicability to increasingly complex genomic contexts and data types. Retriever also has potential applications in several challenging genomic contexts. For polyploid species (e.g., wheat, potato, cotton), which represent many agriculturally important crops, adaptation of the sliding-window approach to accommodate higher ploidy levels could significantly enhance genomic research in these systems. For ancient DNA studies, where highly fragmented and damaged genetic material creates extreme missing data patterns, Retriever’s approach could be combined with damage-aware models to improve recovery of population genetic information from archaeological samples. In clinical genomics, where rare variants often have outsized functional importance, Retriever could be extended to incorporate functional annotation data, prioritizing imputation accuracy in regions with potential phenotypic consequences. For conservation genomics, adaptation to handle extremely small sample sizes through integration with phylogenetic information could enhance genetic management of endangered species with limited sampling opportunities.

As long-read sequencing technologies become increasingly prevalent, extending Retriever to leverage their unique error profiles and capacity for direct haplotype phasing represents another important direction. While the current algorithm is optimized for short-read data characteristics, modifications to the window selection approach could exploit the extended contiguity information provided by long reads, potentially allowing for larger window sizes and more precise boundary definition. The complementary strengths of short-read coverage and long-read contiguity could be combined to further enhance imputation accuracy, particularly for complex structural variants that remain challenging for current approaches.

## Conclusion

Retriever represents a significant advancement in genomic imputation methodology, enabling high-accuracy genotype imputation for non-model organisms without requiring external reference panels. By constructing chimeric reference panels directly from target samples, our approach maintains imputation accuracy exceeding 95% across diverse taxa while preserving essential patterns of linkage disequilibrium. The method demonstrates particular strength in retaining novel variants absent from traditional reference resources. The effectiveness of Retriever rests on the complementary distribution of missing data across samples, the non-random structure of linkage disequilibrium in genomes, and the constrained patterns of allelic variation imposed by population genetic processes. Together, these principles enable the construction of effective reference panels from partially observed data, circumventing the resource-intensive requirements of conventional approaches. Retriever transforms a historical limitation of genomic research into an opportunity, making sophisticated imputation accessible beyond model organism research. In doing so, it contributes to a more inclusive genomic science capable of addressing biological questions across the full diversity of life.

## Author Contributions

D.O-B. conceptualized the chimeric reference panel approach for genomic imputation, developing the hypothesis about the statistical complementarity of missing data patterns across samples. M.Z. implemented this concept by developing the Retriever algorithm, conducting simulations, performing analyses, and creating visualizations. M.Z. received methodological guidance on statistical validation, evolutionary implications, and population genetics from the research team. M.J., J.E. and D.O-B. supervised the project and contributed to experimental design and results interpretation. M.Z. wrote the initial manuscript draft. All authors contributed to manuscript revision, enhancing the theoretical framework, methodology, and discussion to produce the final version.

## Data availability

The source code for Retriever is publicly available through the GitHub repository (https://github.com/uqmzhou8/Retriever.git), released under the MIT license. The repository includes comprehensive documentation, installation instructions, and example scripts to facilitate implementation. All genomic datasets used for validation in this study were obtained from publicly available repositories. Human genomic data were sourced from the 1000 Genomes Project Phase 3 release (http://www.internationalgenome.org/). *Arabidopsis thaliana* sequence data were obtained from the 1001 Genomes Consortium hosted by the Max Planck Institute for Developmental Biology (https://1001genomes.org/). *Gallus gallus* genome sequences were accessed through the study by Cho et al. (2022) available at the European Nucleotide Archive (Project name: PRJEB44919 2021-05-15). *Saccharomyces cerevisiae* genomic data were obtained from Sardi et al. (2018) with accession number PRJEB24747 in EBI.

The specific parameter settings, analytical procedures, and simulated masking patterns used in our evaluation of Retriever are described in detail within the Methods section. Supplementary scripts used for benchmarking, including VCF file processing, performance evaluation, and statistical analysis, are available in the GitHub repository under https://github.com/uqmzhou8/Retriever.git.

Researchers implementing Retriever or reproducing our analyses are requested to cite this paper and the relevant data sources according to their respective citation guidelines. Additional inquiries regarding specific implementations or advanced configurations can be directed to the corresponding author.

## Acknowledgments

We are grateful to members of the Ortiz-Barrientos Laboratory for comments on the figures of this manuscript. We thank Andrew D. Letten for his feedback during the validation process.

## Supporting information

### Comparative analysis of Beagle versions and small sample size considerations

In our methodological evaluation, we observed significant performance difference between Beagle4 and the more recent Beagle5 when applied to small reference panels (Fig S1). This phenomenon warranted deeper investigation, as it directly impacts implementation decisions for non-model organism research where reference sample sizes are inherently constrained.

**Figure S1.**
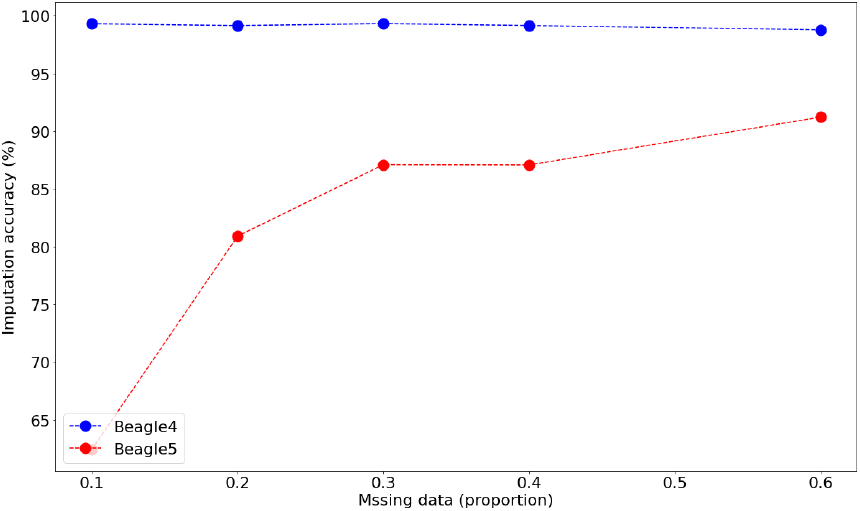
Significant performance difference in the imputation of small external reference panel size (50 individuals) by Beagle4 and Beagle5.

Through correspondence with the principal developer of Beagle5, we received confirmation that the algorithm’s statistical framework was optimized specifically for large-scale human genomic studies with substantial reference panels. The algorithm of Beagle5 implements sophisticated haplotype clustering algorithms that demonstrate substantially improved computational efficiency and imputation accuracy when supplied with reference panels comprising hundreds to thousands of individuals. However, these same algorithmic optimizations create performance instability when the reference panel falls below certain statistical thresholds.

We determined that Beagle4 (v 4.1) maintains robust performance even with modest reference panel sizes (25-50 individuals), making it appropriate for integration with Retriever’s chimeric panel. This finding has significant implications for research communities working with non-model organisms, as it suggests that algorithm selection should be guided by dataset characteristics rather than simply defaulting to the most recent software iteration. Ongoing technical discussions with the Beagle development team aim to characterize the precise statistical thresholds and parameterization adjustments that might enable Beagle5 to function effectively with smaller reference panels. Future releases of Retriever will incorporate these methodological refinements as they become available, potentially extending the computational advantages of newer imputation algorithms to non-model organism research.

### Algorithm details

Retriever constructs the chimeric reference panel by iteratively scanning the genomic positions with a sliding-window. For each window, the algorithm extracts genotype data, removes individuals (columns) with missing genotypes (coded as –3), and then checks whether the number of remaining (complete) individuals meets the user-defined requirement. If the count is insufficient, the window is reduced until the required number is achieved or the minimum window size is reached; if even the smallest window does not yield enough complete data, an error is raised. Otherwise, a random subset of individuals from the complete data is selected to form a reference “bucket”. These buckets, extracted from non-overlapping windows, are then concatenated chronologically to assemble the final chimeric reference panel.

The following pseudocode outlines the method:

#### Algorithm 1 Chimeric Reference Panel Construction

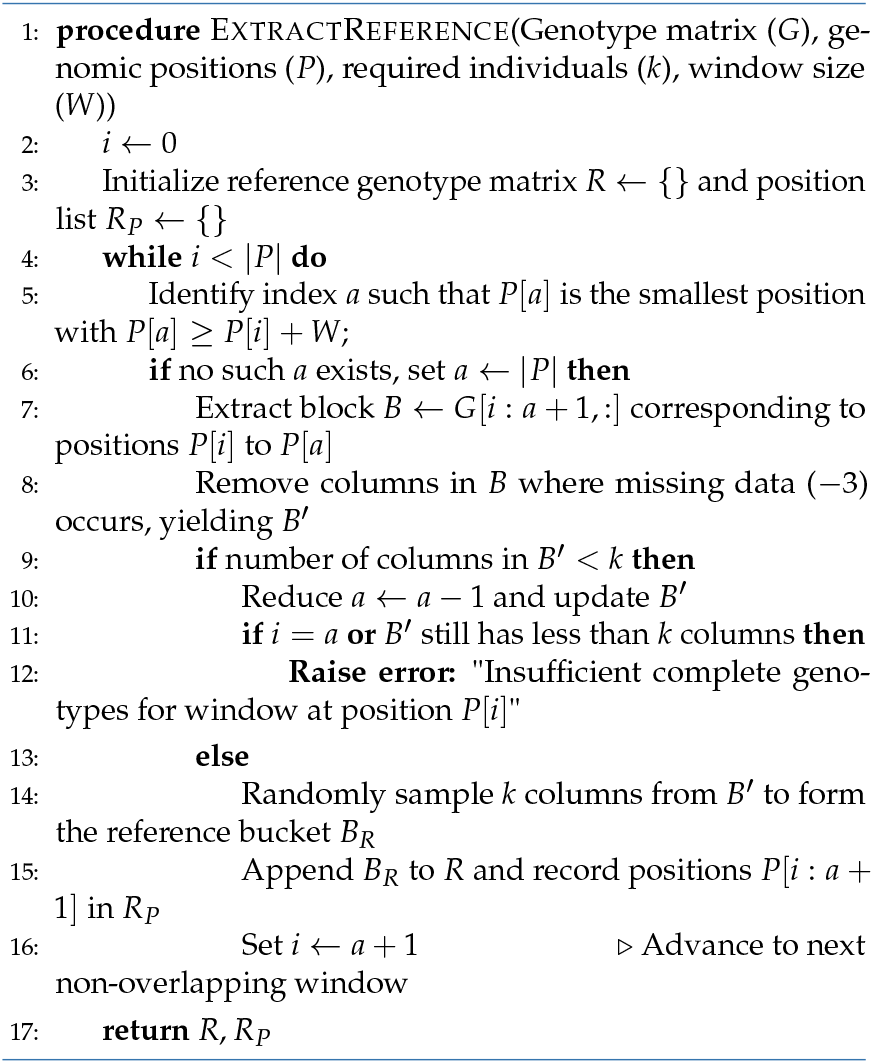

The final chimeric reference panel is obtained by concatenating all buckets *B*_*R*_ extracted across the genome:

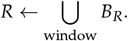

The performance of this approach depends critically on:

- The initial window size *W* (defaulting to 1 kb, or a user-defined value), which determines the genomic span considered in each iteration.
- The minimum window size (e.g. 2 bp), ensuring that data is extracted even in regions with high missingness.
- The required number *k* of individuals per window, which balances the need for sufficient diversity with the constraint of available complete data.

By using this strategy, Retriever leverages the complementary distribution of missing data across samples while preserving the local haplotype structure necessary for effective downstream genotype imputation.

